# Ramping up the Heat: Induction of Systemic and Pulmonary Immune Responses and Metabolic Adaptations in Mice

**DOI:** 10.1101/2025.08.01.667768

**Authors:** Laura E. Dean, Andrea Adamcakova-Dodd, Hans-Joachim Lehmler

**Author notes:** **Corresponding Author:** Dr. Hans-Joachim Lehmler, The University of Iowa, Department of Occupational and Environmental Health University of Iowa Research Park, B164 MTF, Iowa City, IA 52242-5000. **E-Mail Addresses:** (L. E. Dean), (A. Adamcakova-Dodd), and (H. Lehmler).

## Abstract

Heatwaves pose a growing risk to public health. While most animal studies use sudden, extreme heat exposure, the systemic and pulmonary impacts of gradual heat exposures, reflective of real-world conditions, remain poorly characterized. This study examined the effects of acute, gradual extreme heat exposure to mice. Adult male and female C57Bl/6 mice were randomly assigned to heat-exposed, control, or pair-fed groups. Heat-exposed mice experienced a controlled 8-hour temperature ramp from 20°C to 38°C, mimicking the daily transition from nighttime lows to daytime highs. Control and pair-fed mice were maintained in parallel at ambient temperature. Multi-omics profiling was performed to assess cytokine levels in lung and serum, cecal microbiome composition, lung transcriptomics, and serum metabolomics. Heat exposure significantly altered the levels of multiple cytokines in serum and lung, including IL-17α, MIP-1α, MIP-1β, IL-1α, IL-12(p40), and RANTES, indicating shifts in mucosal immunity and immune cell recruitment. Random forest analysis identified 20 taxa that distinguished experimental groups, with a reduction in Lactobacillus observed in males. Lung transcriptomic analysis revealed immune-related gene expression changes involving B cell activation pathways. Serum metabolomics revealed significant decreases in ten metabolites across both sexes, identifying disruptions in amino acid and energy metabolism, with enrichment of the “Glycine, Serine, and Threonine Metabolism” KEGG pathway. Integrative network analyses revealed sex-specific correlations among immune genes, cytokines, and bile acid–related metabolites. These findings show that gradual extreme heat exposure triggers sex-specific systemic and pulmonary immunometabolic responses, offering insight into the biological effects of environmental heat stress and its potential health implications.

**Highlights:**

- Gradual heat exposure altered lung and serum cytokine profiles in mice
- *Lactobacillus* abundance decreased in males, despite stable microbial diversity
- Lung transcriptomics showed B cell–mediated immune activation after heat exposure
- Serum metabolomics revealed heat-induced disruption of amino acid metabolism
- Multi-omics integration revealed sex-specific immunometabolic network responses

## 1. Introduction

Rising global temperatures have increased both the frequency and intensity of extreme heat events (Gao et al., 2023; Marx et al., 2021; Wedler et al., 2023; Yan et al., 2023), posing significant health risks to humans, livestock, and wildlife (Dou et al., 2021; Ebi et al., 2021; Meador et al., 2020; Xiao et al., 2020). Extreme heat is linked to a range of heat-related illnesses (cardiovascular, cerebrovascular, and genitourinary disorders), and a rise in mortality rates, which are anticipated to worsen with ongoing climate change (Daanen & Herweijer, 2015; Ebi et al., 2021; Folkerts et al., 2020; Périard et al., 2016). Furthermore, exposure to extreme heat can trigger systemic inflammation that affects both the brain and peripheral organs, ultimately leading to systemic inflammatory response syndrome (SIRS) (Leon & Helwig, 2010). Heatwaves also increase morbidity and mortality in patients with chronic lung diseases, such as chronic obstructive pulmonary disease (COPD) (Konstantinoudis et al., 2022; Scheerens et al., 2022; Silveira et al., 2023; Witt et al., 2015), and exacerbate the adverse effects of air pollution (Grigorieva & Lukyanets, 2021). Therefore, major health organizations, including the World Health Organization, have prioritized further research into the health impacts of extreme heat to develop effective adaptation and mitigation strategies (Bernstein & Rice, 2013; Crowley et al., 2016; Huang et al., 2015; McCormack et al., 2016; Patz et al., 2014; Pinkerton et al., 2012; Portier et al., 2010; World Health Organization, 2012).

Mammalian thermoregulation during acute heat exposure involves rapid physiological responses, such as vasodilation, increased heart rate to facilitate heat transfer to the skin, and enhanced sweat production for evaporative cooling (Corbett et al., 2018; Ebi et al., 2021; Navas-Martín et al., 2024). Acute heat can also compromise respiratory function, disrupt endothelial barriers, and trigger systemic inflammatory responses that culminate in widespread tissue damage, leading to heat stress, heat shock, and, ultimately, death (Ebi et al., 2021; Lepeule et al., 2018; Miao et al., 2022). During heatwaves, the repeated activation of the heat shock response and subsequent increase in heat shock protein expression can lead to improved cellular protection against heat stress (Sartori & Scherrer, 2003; Wheeler & Wong, 2007). This cellular-level adaptation complements the systemic physiological changes, such as improved sweating and cardiovascular responses, that enhance overall heat tolerance (Racinais et al., 2019).

The effects of sudden, extreme heat exposure have been extensively investigated in animal models, such as rats (Chauhan et al., 2017; Dou et al., 2021), mice (Lee et al., 2015; Leon et al., 2005; Miyamoto et al., 2021; Roths et al., 2024), or pigs (Lebret et al., 2023; Mayorga et al., 2020; Ross et al., 2017). Similarly, laboratory studies of human responses to extreme heat use sudden heat exposure paradigms (Meade et al., 2024). These experimental paradigms simulate scenarios such as stepping from an air-conditioned space into a hot outdoor environment. However, few studies have examined the consequences of a gradual increase in outdoor or indoor temperature that reflects the biological responses to real-world conditions (Ahrens, 2009). In outdoor settings, gradual heat exposures are, for example, experienced by farmers (Elliott et al., 2022), athletes (Sawka et al., 1993), and military personnel (Alele et al., 2020; Parsons et al., 2019). Importantly, populations vulnerable to extreme heat, such as children and elderly individuals (Basu, 2009; D’Amato & D’Amato, 2023; Leon & Helwig, 2010; Lepeule et al., 2018), face elevated risks when residing in buildings without adequate air conditioning (Baniassadi et al., 2019), with evidence suggesting that indoor temperatures during extreme heat events correlate to outdoor temperatures (Nguyen et al., 2014).

In this study, we employ a systems biology approach to investigate the pulmonary and systemic effects of gradual extreme heat exposure in mice. We comprehensively assess inflammatory markers in serum and the lung, cecal microbiota composition, lung gene expression, and serum metabolite profiles to elucidate how a gradual increase in temperature influences these physiological and molecular parameters. Our findings suggest that gradual extreme heat exposure, hereafter referred to as heat exposure, triggers sex-specific systemic and pulmonary immune and metabolic responses.

## 2. Experimental

### 2.1. Animals and exposure

All experimental procedures involving animals were reviewed and approved in accordance with all Public Health Service policies and the Guide for the Care and Use of Laboratory Animals, NIH Publication No. 85-23, revised 2011, by the University of Iowa IACUC (Institutional Animal Care and Use Committee) and conform with the National Research Council’s Guide for the Care and Use of Laboratory Animals (National Research Council, 1981). Details regarding the animal study are reported in accordance with the ARRIVE guidelines.

Animals were housed in an animal facility at the University of Iowa, accredited by the Association for Assessment and Accreditation of Laboratory Animal Care International (AAALAC), in a temperature and humidity-controlled room (23°C, 55% relative humidity) with a 12 h light/dark cycle before heat exposure.

Adult, 10-week-old C57Bl/6 mice (n = 24 per sex) were obtained from Jackson Labs (Bar Harbor, ME, USA). At the start of the experiment, male (25.5 ± 1.3 g) and female (20.4 ± 1.3 g) mice were randomly assigned to one of three experimental groups (heat-exposed, control, or pair-fed) using the random number generator function of Microsoft Excel (version 2308). Female mice were estrus synchronized using 5 IU of pregnant mare serum (Prospec, East Brunswick, NJ, USA) and 5 IU of human chorionic gonadotropin (Sigma Aldrich, St. Louis, MO, USA). This synchronization occurred five days and three days, respectively, before exposure to the heat event. The purpose of this procedure was to ensure that all females were at the same stage of their reproductive cycle during the heat exposure, thus eliminating the estrous cycle as a potential confounding factor (Ajayi & Akhigbe, 2020; Goldman et al., 2007). All animals had access to Teklad-7913 rodent chow (ENVIGO, Indianapolis, IN, USA) and water *ad libitum* throughout the experiment, except for the pair-fed mice. Pair-fed mice had access to the same amount of food and water that the heat-exposed mice had consumed during the 8-hour heat event to control for differences in food and water intake. This control group enabled us to determine whether the observed effects are due to heat exposure or variations in diet and hydration. No adverse effects were observed during the study.

Mice, after their arrival, were acclimated for two (males) or three (females) weeks to the animal facility (including new water and food dispensers) in the Metabolic Phenotyping Core Facility at the University of Iowa. Female mice were given an extra week of acclimation due to the difference in body weights and size between males and females. Bedding was mixed to synchronize the microbiome before heat exposure twice a week throughout the acclimation period (Miyoshi et al., 2018; Van Loo et al., 2000). Heat-exposed mice were placed into an incubator (Powers Scientific, Inc., Doylestown, PA, USA) and the temperature was gradually increased from 20°C to 38°C. The control and pair-fed mice were placed on a benchtop in the same room as the incubator and maintained at an ambient temperature of 23°C for 8 hours. The incubator humidity was 61 ± 8%, and the room humidity was 55 ± 1%.

All animals were euthanized via cervical dislocation and decapitation immediately after 8 hours of exposure. Trunk blood was collected, and the serum was prepared and stored at −80°C for further analyses. Lungs were flash frozen and stored at −80°C for cytokine analysis or placed in RNAlater buffer and stored at −20°C for RNA sequencing. Cecum content was collected, flash frozen to determine changes in the microbiome, and stored at −80 °C for later analysis.

### 2.2. Cytokine analysis

Lung tissues were homogenized in T-PER buffer (Thermo Scientific, Waltham, MA, USA) at a ratio of 1 mg of tissue to 20 μL of buffer for cytokine and chemokine measurements. Homogenates were centrifuged at 10,000 ×g for 5 minutes at 4°C, and the resulting supernatant was collected. Total protein concentrations were determined using the Pierce BCA Protein Assay Kit (catalog #23225, Thermo Scientific) following the manufacturer’s instructions.

Cytokine and chemokine levels, including IL-1α, IL-1β, IL-2, IL-3, IL-4, IL-5, IL-6, IL-9, IL-10, IL-12 (p40), IL-12 (p70), IL-13, IL-17A, Eotaxin, G-CSF, GM-CSF, IFN-γ, KC, MCP-1 (MCAF), MIP-1α, MIP-1β, RANTES, and TNF-α (**Table S1**), were quantified in both serum and lung homogenates using the Bio-Plex Pro Mouse Cytokine 23-plex assay (Bio-Rad Laboratories) according to the manufacturer’s protocol. A Luminex 200 system (Bio-Rad Laboratories) was employed for data acquisition and analysis, as previously described (Areecheewakul et al., 2023). Serial dilutions of the lyophilized cytokine standard were made using the assay diluent provided with the kit. For lung homogenates, a 1:1 mixture of assay diluent and T-PER reagent was used. Serum samples were diluted 1:4, while lung homogenates were diluted 1:5 in their respective diluents before analysis.

### 2.3. Microbiome analysis

DNA was extracted from each cecum sample using the DNeasy PowerLyzer PowerSoil Kit (Qiagen, Germantown, MD, USA) following the manufacturer’s instructions. DNA levels were quantified using a Nanodrop (Thermo Scientific) and stored at −80°C until further analysis. Sequencing was performed on the Illumina MiSeq platform (Illumina, San Diego, CA, USA) by amplifying the V3-V4 region of the 16S rRNA genes. Downstream analyses of fastq files were conducted using the dada2 script in R (Callahan et al., 2016; Shahi et al., 2020) to generate amplicon sequence variants (ASVs). The ASV table was analyzed using MicrobiomeAnalyst (Chong et al., 2020; Dhariwal et al., 2017). Features were filtered based on the inter-quantile range, with a minimum count of 200, a prevalence in the sample equal to 20%, and a removal percentage of 10%. No rarefication or scaling was applied to normalize the data. However, the data was transformed using relative log expression before subsequent analyses. Alpha diversity, a common measurement of microbial diversity within an experimental group, was assessed at the feature level using Shannon diversity (Kers & Saccenti, 2022). Beta diversity, a measure of microbial diversity between experimental groups, was measured using Principal Coordinates Analysis (PCoA) and Jensen-Shannon Divergence (Chen et al., 2021; Kers & Saccenti, 2022). Additionally, random forest was employed to identify differentially abundant taxa (5000 trees and 7 predictors, with randomness enabled). Subsequently, phylogenetic investigation of communities by reconstruction of unobserved states (PICRUSt) was employed to predict the functional potential of the microbial communities (Langille et al., 2013).

### 2.4. RNA sequencing and analysis

Total RNA was isolated using the RNeasy Mini Kit (Qiagen) from the hippocampus and left lung lobe following the manufacturer’s instructions. RNA was checked for purity with a nanodrop (Thermo Fisher Scientific, Fair Lawn, NJ, USA). A total of 158 samples with RNA integrity values of ≥ 8.0 were submitted to Novogene (Sacramento, CA, USA) for RNA sequencing using the NovaSeq 6000 PE150 platform. Novogene conducted an additional quality control step using an Agilent 5400 Fragment Analyzer System (Agilent Technologies, Santa Clara, CA, USA), followed by cDNA library preparation employing ribosomal depletion techniques. Sequencing was then carried out to a depth of 20 million reads, with 120 base-pair paired-end sequences following the Illumina 1.9 encoding protocols established by Novogene.

Raw RNA sequencing FASTQ files were generated by Novogene through demultiplexing and concatenation for each sample. Further processing for differential gene expression was done based on methods previously described (Bullert et al., 2024). Briefly, gene counts were determined by *GenomicAlignments* (R and Rstudio version 4.2.2) using Novogene’s reference genome. Differential expression analysis was then performed using DESeq2 (version 1.38.3) (Love et al., 2014) and visualized through principal component analysis (PCA) plots using the first two principal components and volcano plots using *EnhancedVolcano* (Blighe et al., 2024). Because >30 differentially expressed genes (DEGs) were observed in heat-exposed mice, further analysis of pathways was conducted for further interpretation of the data via iPathwayGuide (Advaita Corporation, Ann Arbor, Michigan, USA) as described previously (Bullert et al., 2024).

### 2.5. Metabolomics LC-MS instrument parameters

Serum metabolomics analyses were performed at the Northwest Metabolomics Research Center (University of Washington, Seattle, WA, USA) with an AB Sciex 6500+ Triple Quadrupole MS (Sciex, Framingham, MA, USA) with an ESI ionization source and equipped with two Shimadzu UPLC pumps (Shimadzu Corporation, Kyoto, Japan) and a CTC Analytics PAL HTC-xt temperature-controlled auto-sampler (CTC Analytics AG, Zwingen, Switzerland) (Meador et al., 2020). Each sample was injected into two identical hydrophilic interaction liquid chromatography (HILIC) columns (Waters XBridge BEH Amide XP, Waters Corporation, Milford, MA, USA) on the liquid chromatography tandem mass spectrometry (LC-MS/MS) system. One column operated in ESI+ mode, and the other in ESI-mode, with mass spectrometry data acquired in the multiple-reaction monitoring (MRM) mode. The LC-MS/MS system was operated using AB Sciex Analyst 1.6.3 software, while the chromatographic peaks were integrated with AB Sciex MultiQuant 3.0.3 software. The analytical method targeted 361 metabolites and 32 stable isotope-labeled internal standards (SILISs). In this study, 185 metabolites and 30 SILISs were detected. We employed two quality control (QC) sample types: pooled human serum samples to monitor system performance and pooled study samples to assess data consistency. These QC samples were analyzed after every 10 experimental samples and demonstrated consistency with an average coefficient of variation (CV) of 2.2%.

### 2.6. Metabolomics data analysis and visualization

Metabolomics data were analyzed with the MetaboAnalyst 6.0 online platform (Pang et al., 2021). Briefly, all data were normalized by sum and log-transformed. Differences between experimental groups were analyzed using Partial Least Squares Discriminant Analysis (PLS-DA), random forest modeling, and Kyoto Encyclopedia of Genes and Genomes (KEGG) pathway analysis.

### 2.7. Correlation analysis

Integrated network analyses were performed for male and female mice using xMWAS (v. 1.0) (Uppal et al., 2018). The goal of these analyses was to explore the relationships between lung gene expression, serum metabolites, serum cytokines, and lung cytokines across exposure groups (heat-exposed vs. control, heat-exposed vs. pair-fed, and pair-fed vs. control). xMWAS calculates pairwise association scores, which approximate correlation coefficients, between each cytokine level, metabolite level, or gene expression. This was accomplished using partial least squares (PLS) regression analysis. xMWAS then generated a multidata integrative network (Cao et al., 2018). Only associations with specified correlation threshold (R) and p-value (determined by Student’s t-test) were included in this network (R > 0.9 and p-value < 0.05). Finally, a multilevel community detection algorithm was applied (Blondel et al., 2008) to identify clusters genes, metabolites, and cytokines that exhibited strong internal connections but sparse external connections. Cytoscape (v. 3.10.1) was used to visualize the correlation networks from the xMWAS analyses (Otasek et al., 2019; Shannon et al., 2003).

### 2.8 Statistical analysis

Body and tissue weights are reported as mean ± standard deviation. Comparisons between all groups for each respective cytokine were performed by a 2-way ANOVA with Tukey multiple comparisons using GraphPad Prism (GraphPad Software, Boston, MA, USA; RRID:SCR_002798). Log fold change and p-values were calculated based on a Mann Whitney U test of the ASV data for features that were statistically significant according to random forest analysis (mean decrease in accuracy (MDA) > 0.01) and PICRUSt analysis.

## 3. Results and Discussion

### 3.1. General responses to heat exposure

Body weights and organ weights were not significantly different between experimental groups (**Figure 1A, Table S2**). Feed consumption tended to increase in heat-exposed mice compared to both control and pair-fed male and female mice; however, this difference did not reach statistical significance (**Figure 1B**). Unlike prior studies of sudden heat exposure, where reduced feed intake in heat-exposed mice mitigated the thermal effect of digestion, we hypothesize that our gradual heat exposure protocol may have elicited a different behavioral adaptation, which could explaining the difference between our findings and those of others (Baumgard & Rhoads, 2013; Morera et al., 2012; Roach et al., 2024; Xiao et al., 2020).

**Figure 1.**
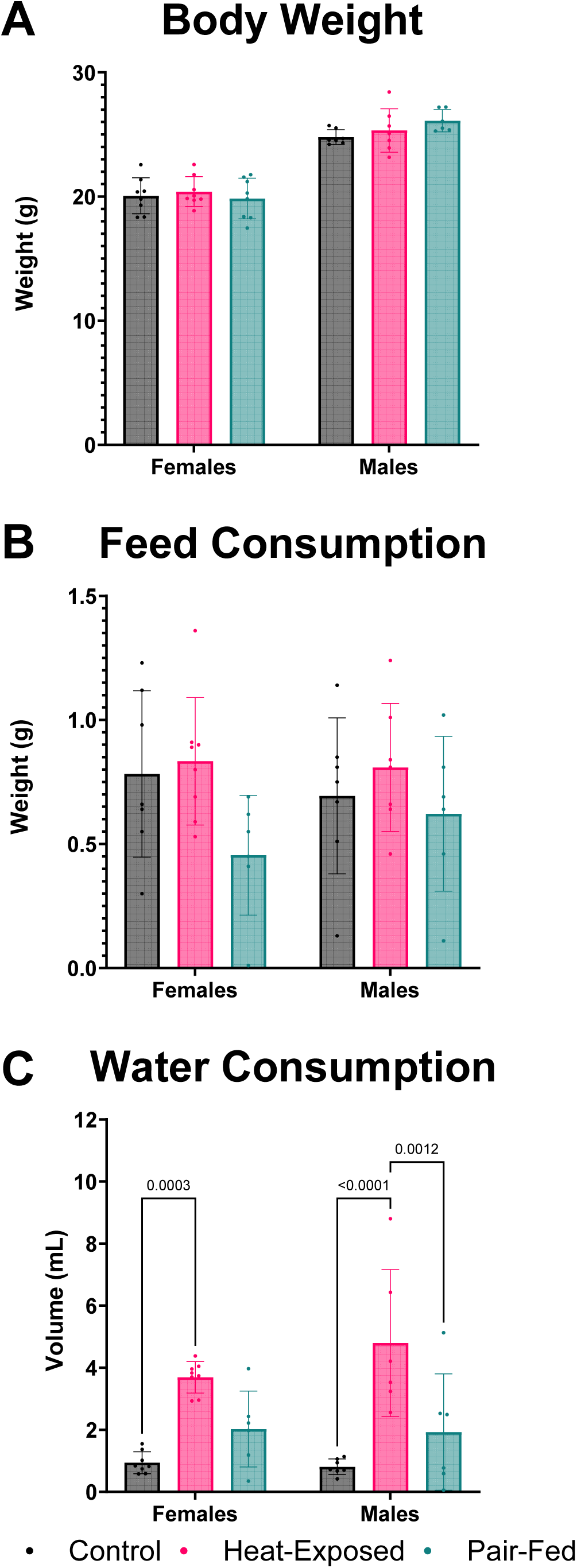

Consistent with our expectations, heat-exposed mice exhibited significantly higher water consumption compared to their control and pair-fed counterparts (**Figure 1C**). This increase is attributed to the increased water intake in mammals to maintain thermal balance during elevated temperatures, thereby supporting physiological adaptation (National Research Council, 1981; Wen et al., 2021).

### 3.2. Cytokine and chemokine alterations following heat exposure

Exposure to heat can trigger systemic inflammation and disrupt immune function, as evidenced by increased circulating cytokine and chemokine levels (Cantet et al., 2021; Chauhan et al., 2017; Di Vincenzo et al., 2024). Therefore, we assessed cytokine and chemokine levels in lung tissue and serum to characterize pulmonary and systemic inflammatory responses induced by heat exposure.

#### 3.2.1 Lung cytokine and chemokine levels

Heat exposure significantly affected 3 out of 23 measured cytokines and chemokines in lung homogenates (**Figure 2A-C**). In male mice, IL-17α levels were significantly lower in the heat-exposed group compared to the pair-fed group and exhibited a decreasing trend compared to the controls. In female mice, IL-17α levels were not significantly altered by heat exposure or pair-feeding; however, a trend toward decreased levels was observed in both heat-exposed and pair-fed groups compared to the control group. Conversely, MIP-1α (CCL3) levels were significantly increased in female mice exposed to heat compared to both the control and pair-fed groups. MIP-1β (CCL4) levels in females were significantly elevated in the heat-exposed group compared to the pair-fed group, with a similar trend observed compared to the control group. In male mice, neither MIP-1α nor MIP-1β levels were significantly altered by heat exposure, though both exhibited an increasing trend in the heat-exposed group relative to control and pair-fed groups.

**Figure 2.**
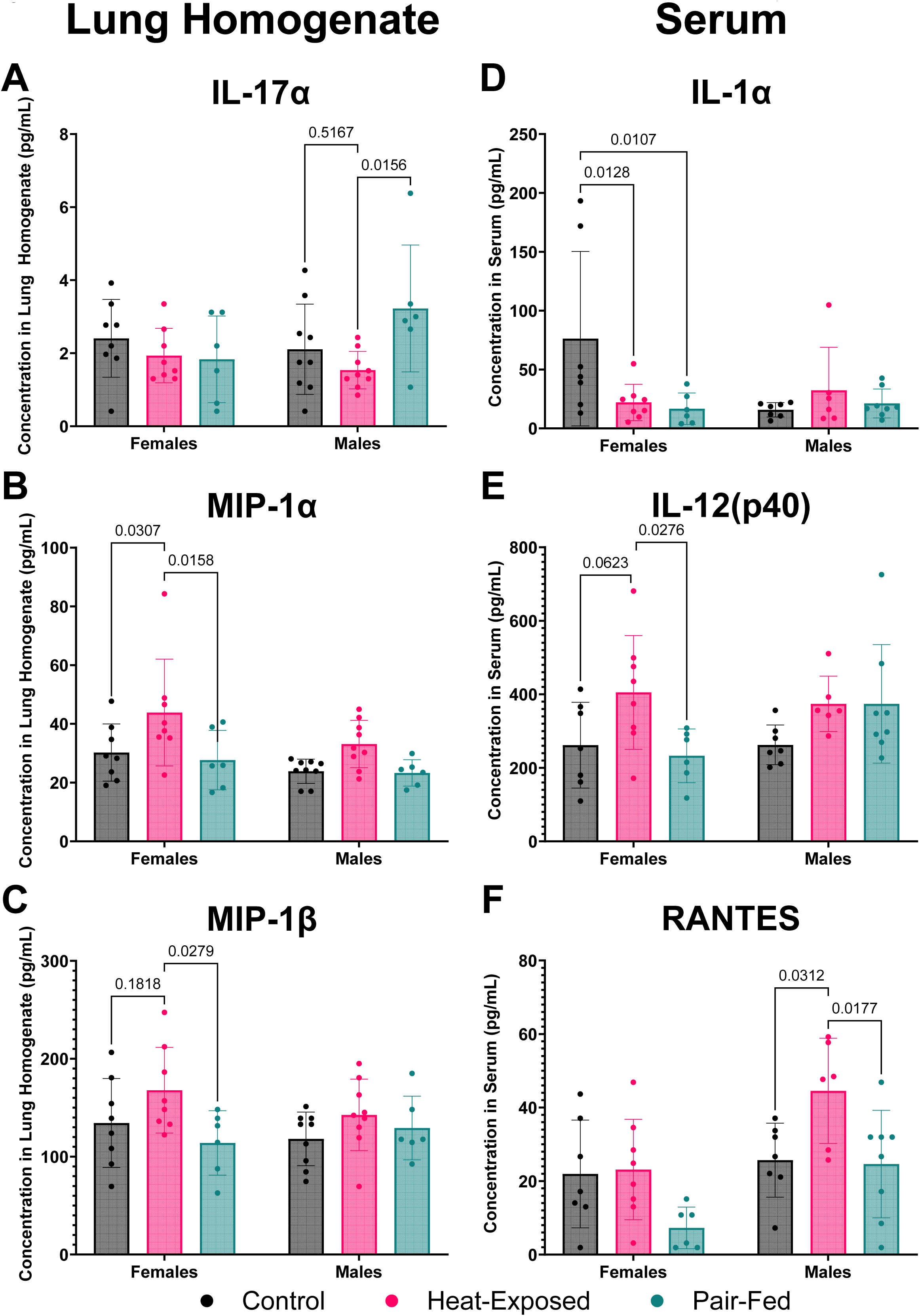

The reduction of IL-17α in males and the elevation of MIP-1α and MIP-1β in both sexes in heat-exposed mice are caused by heat exposure rather than reduced feed intake. The decrease in IL-17α in the lung may indicate a potential impairment in mucosal defense, as IL-17α plays a crucial role in maintaining mucosal immunity (Gurczynski & Moore, 2018). Changes in IL-17α levels also align with previous research demonstrating that IL-17α plays a role in maintaining health during the response to injury, physiological stress, or infection (McGeachy et al., 2019). Elevated levels of MIP-1α and MIP-1β in the lungs suggest increased immune cell recruitment and activation, indicative of an inflammatory response to heat stress (Lillard et al., 2003). In contrast, the alterations in IL-17α levels in heat-exposed and pair-fed female mice compared to controls likely indicate an effect related to decreased feed intake.

#### 3.2.2. Serum cytokine and chemokine levels

Heat exposure significantly altered 3 out of the 23 measured cytokines and chemokines in serum (**Figure 2D-F**). IL-1α levels were significantly decreased in heat-exposed and pair-fed female mice compared to the control group but were unaffected in male mice. IL-12(p40) was significantly elevated in heat-exposed female mice compared to the pair-fed group and showed a trend toward increased levels compared to controls. In male mice, IL-12(p40) also showed a trend toward elevation in both the heat-exposed and pair-fed groups compared to the control group. RANTES (CCL5) levels were significantly increased in heat-exposed male mice compared to both the control and pair-fed groups, a trend that was also observed in female mice.

The observed alterations in IL-1α in male mice, IL-12(p40) in females, and RANTES in both sexes suggest that these changes are driven by heat exposure rather than feed intake. In contrast, the changes in IL-1α in heat-exposed and pair-fed female mice and IL-12(p40) in heat-exposed and pair-fed male mice suggest that reduced feed intake contributes to some of these effects. These heat-specific effects on serum cytokine and chemokine levels suggest a shift in the immune response under heat exposure. The reduction in IL-1α in the serum of male mice, a cytokine associated with acute inflammation, may indicate a shift away from an early pro-inflammatory response (Dinarello, 2018). Meanwhile, the elevation of IL-12(p40) in the serum of heat-exposed female mice indicates a strong immune response, driving Th1 responses (Trinchieri, 2003). Increased levels of RANTES (CCL5) in the serum of male mice and a similar trend in female mice suggest heightened immune cell recruitment, a hallmark of chronic inflammation or immune activation (Ajuebor et al., 2001; Koya et al., 2006; Machura et al., 2024; Zeng et al., 2022). Interestingly, chemokines that were increased after the heat exposure either in lungs and serum such as MIP-1α (CCL3), MIP-1β (CCL4), and RANTES (CCL5) belong to the CC sub-family of chemokines, which are known for their role in leukocyte trafficking, recruitment and activation (Zeng et al., 2022). Since other pro-inflammatory cytokines and chemokines measured in serum were not significantly altered by acute heat exposure, we are hesitant to conclude that there was a robust effect on systemic inflammation in this study. However, if exposure to heat is repeated and the inflammation is not completely resolved before the next heat insult, it may lead to chronic inflammation. Moreover, RNA sequencing results show the activation of immune responses, especially in the lungs, as discussed below.

### 3.3. Microbiome changes following heat exposure

Changes in the cecal microbiota were examined after heat exposure, as the microbiome influences inflammatory markers throughout the body (Di Vincenzo et al., 2024; Haase et al., 2018; Wen & Duffy, 2017). Moreover, previous studies have shown that sudden changes in environmental temperature modify gut microbiome composition and function as well as the intestinal barrier (Hylander & Repasky, 2019; Wang et al., 2024c; Wen et al., 2021; Zhou et al., 2024). Unlike the earlier studies, we observed no significant changes in alpha diversity in female or male mice (**Figure S1A-B**). Heat exposure also did not affect the beta diversity of the microbial populations compared to control or pair-fed female or male mice (**Figure S1C-D**). A random forest analysis identified 20 taxa (comprising four phyla, five classes, five orders, three families, two genera, and one species) that were key in differentiating the experimental groups (**Figure 3**). However, a Mann-Whitney U test indicated that the relative abundances of these taxa were not significantly affected by heat exposure or controlled feed intake. Despite this, many taxa identified through random forest analysis exhibited a sex-dependent decrease in abundance based on fold change following heat exposure. For example, the genus *Lactobacillus* was reduced in male mice exposed to heat, while its levels remained unchanged in female mice. The reduction in *Lactobacillus* abundance may indicate compromised gut health following exposure to the gradual extreme heat event, as these bacteria are essential for maintaining mucosal integrity and modulating immune responses (Rastogi & Singh, 2022; Wells, 2011).

**Figure 3.**
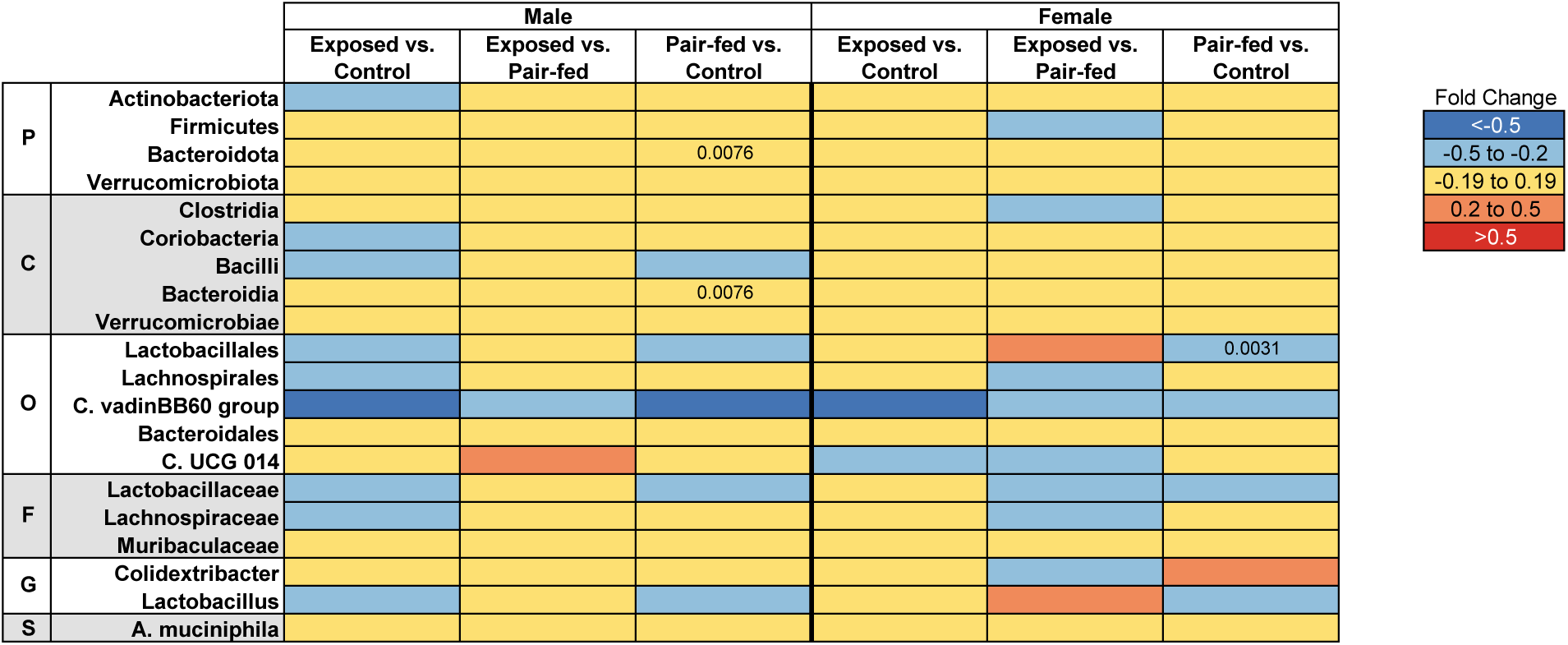

### 3.4. Transcriptomic effects in lungs and hippocampus after heat exposure

Sudden heat exposure (heat stress) is a known cause of transcriptomic responses in the lung (Liu et al., 2020; Nan et al., 2023; Wang et al., 2024b). Additionally, heat stress has been shown to lead to significant changes in the expression of gene expression implicated in heat stress-induced cognitive impairment in the hippocampus (Li et al., 2025; von Ziegler et al., 2022). Considering that the central nervous system regulates a wide range of bodily functions and previous studies found transcriptomic changes in the lung, we performed RNA sequencing of both the hippocampus and the lung.

The PCA did not reveal a separation of gene expression profiles in the lungs or hippocampus of male or female mice by experimental groups (**Figure S2**). We also did not identify any significant DEGs in the hippocampus. However, we observed significant transcriptomic alterations in the lung following heat exposure. Therefore, iPathwayGuide was performed to characterize biological processes affected by heat exposure in the lung. Heat exposure mainly impacted biological processes related to the activation of immune responses, consistent with the cytokine and chemokine data discussed above, in comparison to the control and pair-fed groups (**Figures 4** and **5**). This observation supports the previously documented role of the lung in heat-induced immune responses (Lepeule et al., 2018; Wheeler & Wong, 2007). Notably, though not observed in iPathway Guide analysis, the observed changes mirrored inflammatory responses seen in sepsis-induced acute respiratory distress syndrome (ARDS) (Bouchama & Knochel, 2002; Cheng-Kuei et al., 2006; Yang et al., 2012).

#### 3.4.1. Analysis of biological processes altered in lungs in heat-exposed vs. control groups

In female mice, heat exposure led to the upregulation of 90 genes and the downregulation of 31 genes when compared to controls (**Figure 4A**). Similarly, in male mice, 105 genes were upregulated and 48 were downregulated relative to controls (**Figure 5A**). In the iPathwayGuide analysis, two immune-related biological processes, “Immune System Process” and “Lymphocyte Activation,” were significantly altered in both heat-exposed female and male mice compared to controls (**Figures 4C** and **5C**). “Immunoglobulin-Mediated Immune Response,” “B Cell-Mediated Immunity,” and “Immune Response” were altered only in heat-exposed female mice compared to controls (**Figure 4C**). Twenty-two genes were associated with these biological processes, with the expression of 10 genes upregulated and 12 genes downregulated in female mice following heat exposure. In male mice, heat exposure affected additional immune-related processes, for example, “Leukocyte Activation,” “Cell Activation,” and “B Cell Activation.” Sixteen genes were associated with these biological processes, with the expression of 13 genes significantly upregulated, and three downregulated in heat-exposed male mice.

**Figure 4.**
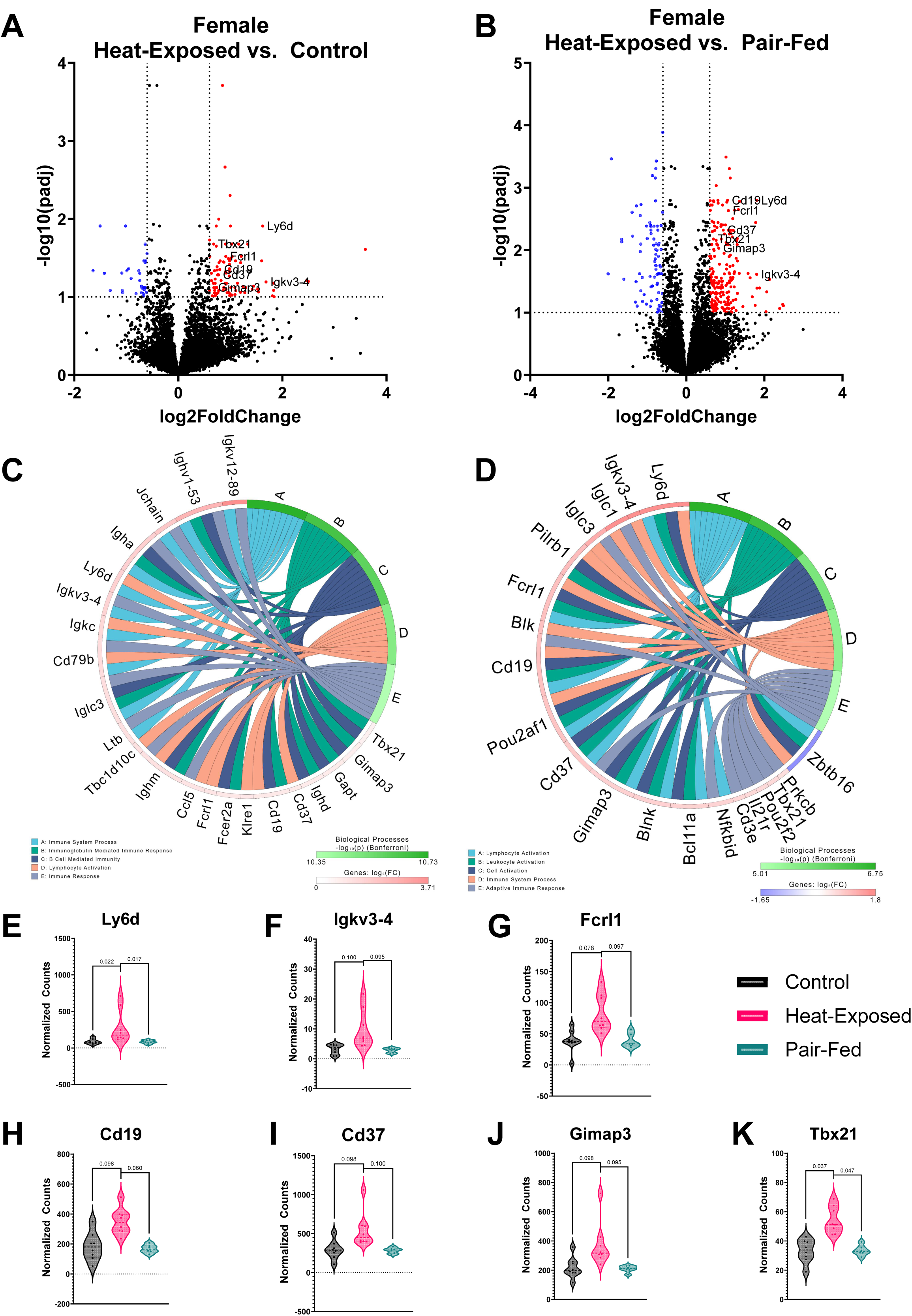

Four genes were upregulated in both male and female heat-exposed mice compared to controls, including Fc fragment of IgE receptor II (*Fcer2a*), CD19 molecule (*Cd19*), Immunoglobulin heavy constant delta (*Ighd*), and GRB2-binding adaptor protein, transmembrane (*Gapt*) (**Figure S3**). *Fcer2a* was associated with “Immunoglobulin Mediated Immune Response” and “B Cell-Mediated Immunity” in females and “Leukocyte Activation,” “Cell Activation,” and “Immune System Process” in males. *Fcer2a* modulates immune responses in the lung and is expressed on various immune cells within the lung, including B cells and macrophages (Cernadas et al., 1999). *Cd19*, which positively regulates B cell activation (Karnell et al., 2014), was associated with “Immunoglobulin Mediated Immune Response,” “B Cell-Mediated Immunity,” and “Lymphocyte Activation” in females and the top five biological processes in males. *Ighd*, which is involved in B cell receptor signaling and adaptive immune response (Schroeder & Cavacini, 2010), was associated with “Immunoglobulin Mediated Immune Response” and “B Cell Mediated Immunity” in female and “B Cell Activation” in male mice. Finally, *Gapt*, which plays an important role in B cell activation (Liu & Zhang, 2008), was associated with “Immunoglobulin Mediated Immune Response” and “B Cell-Mediated Immunity” in females and the top five biological processes in males. Overall, these gene expression changes in heat-exposed mouse lungs signal a B-cell-mediated component of the gradual heat stress response (Cernadas et al., 1999; Karnell et al., 2014; Liu & Zhang, 2008; Schroeder & Cavacini, 2010).

#### 3.4.2. Analysis of biological processes altered in lungs in heat-exposed vs. pair-fed groups

Compared to pair-fed animals, heat exposure resulted in the upregulation of 91 genes and downregulation of 46 genes in females (Figure 4B) and the upregulation of 207 genes and downregulation of 80 genes in males (Figure 5B). In the iPathwayGuide analysis, several immune-related biological processes, including “Immune System Process,” “Cell Activation,” “Leukocyte Activation,” and “Lymphocyte Activation,” were significantly altered in heat-exposed female and male mice compared to pair-fed animals (**Figures 4D** and **5D**). In addition, “Immune System Process” and “Adaptive Immune Response” were altered in heat-exposed vs. pair-fed female mice (**Figure 4D**), with 18 significantly elevated and one significantly decreased gene associated with these biological processes. In male heat-exposed vs. pair-fed mice, “Cell Activation” and “Immune Response” were altered (**Figure 5D**). There were 15 significantly elevated and two significantly decreased genes associated with these biological processes in heat-exposed vs. pair-fed male mice. No shared DEGs were detected between males and females, despite overlapping biological processes.

**Figure 5.**
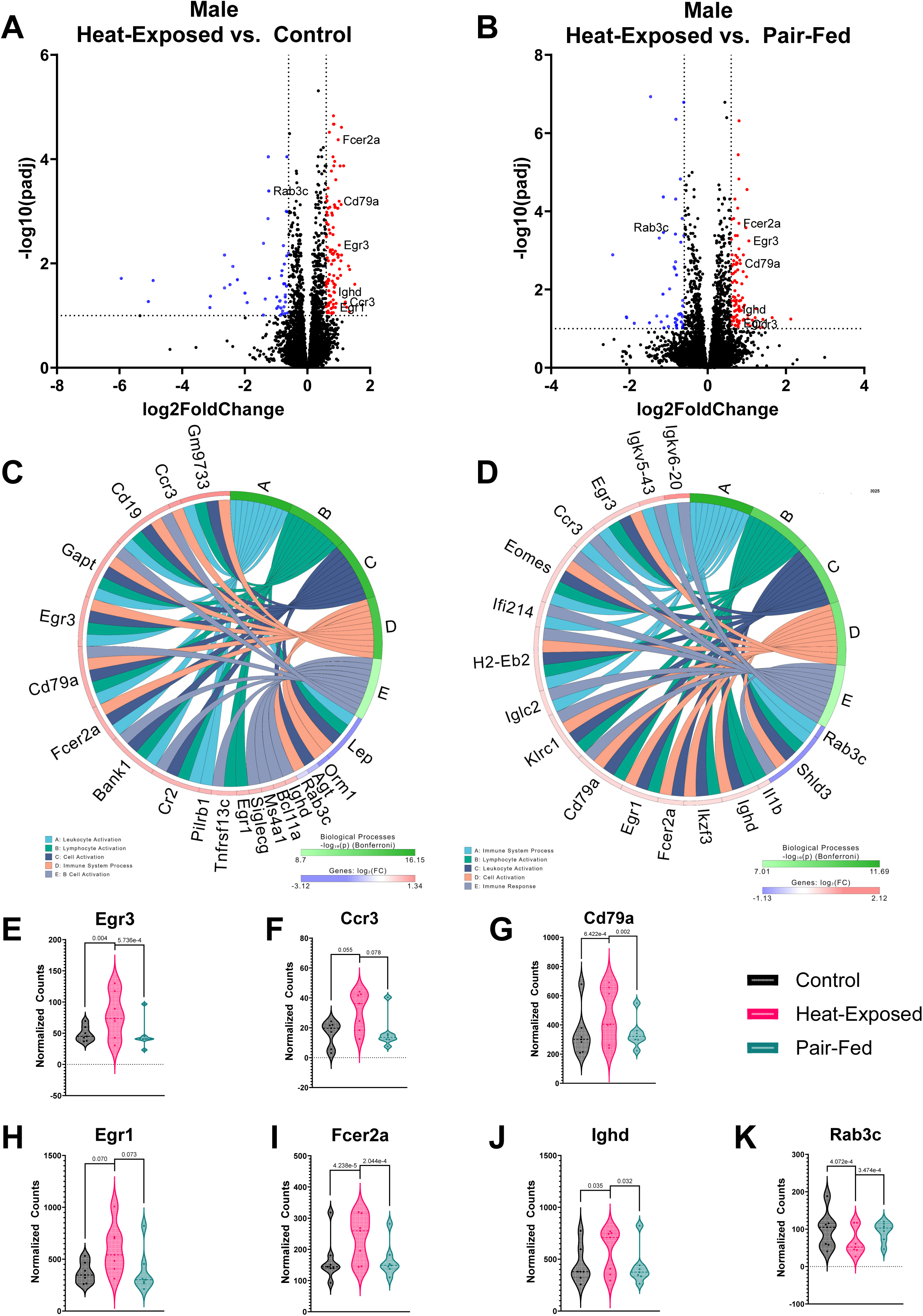

#### 3.4.3. Shared lung transcriptomic responses to heat exposure

Heat exposure led to alterations in two biological processes, “Immune System Process” and “Lymphocyte Activation,” compared to the control and pair-fed groups in females. This was associated with seven shared DEGs: *Ly6d, Igkv3-4, Fcrl1, Cd19, Cd37, Gimap3,* and *Tbx21* (**Figures 4E-K**). In males, heat exposure altered four biological processes, “Immune System Process,” “Lymphocyte Activation,” “Leukocyte Activation,” and “Cell Activation,” compared to both the control and pair-fed groups, with seven shared DEGs: *Egr3, Ccr3, Cd79a, Egr1, Fcer2a, Ighd,* and *Rab3c* (**Figures 5E-K**). This 8-hour heat exposure affected multiple immune-related genes, particularly those involved in B-cell activation (*CD19, CD79a, CD37, FCRL1*), immunoglobulin production (*Igkv3-4, Ighd*), and inflammatory signaling (*Egr1, Rab3c*), in the mouse lung (Morbach et al., 2016; Zou & Zeng, 2023). Transcription factors like *Tbx21* (T-bet) and *Egr3* regulate immune responses under heat stress, while chemokine receptors (*CCR3*) and survival-associated genes (*Gimap3*) contribute to immune cell recruitment and persistence in the lung (Conroy & Williams, 2001; Huang & Bi, 2021). Collectively, these results reveal a heat-induced, B cell-centric immune activation in the lung, with partially overlapping sex-specific molecular signatures.

### 3.5. Serum metabolome alterations following heat exposure

Serum metabolomics was performed to identify potential biomarkers related to heat exposure. Metabolomics analyses revealed significant alterations in endogenous metabolites in both male and female mice compared to their respective control and pair-fed groups. PLS-DA plots of females and males showed a separation between the heat-exposed and control groups, with the pair-fed group overlapping the other two experimental groups (**Figures 6A** and **7A**). Heatmaps illustrating the top 25 metabolites affected by heat exposure show that 10 metabolites were altered in both female and male mice. These metabolites include methionine sulfoxide, 4-hydroxybenzoic acid, leucine, valine, isoleucine, isovalerylcarnitine, dihydrouracil, ornithine, threonine, and sarcosine (**Figures 6B** and **7B**). All 10 metabolites decreased in abundance following heat exposure compared to both control and pair-fed groups.

#### 3.5.1. Random forest analysis of metabolites

Based on random forest analyses, several metabolites were important in distinguishing the metabolomes of the experimental groups. In females, N-Ac-glutamate, pseudouridine, and N-Ac-alanine had MDA scores > 0.01 (**Figure 6C**). N-acetyl-glutamate was decreased in heat-exposed mice, while pseudouridine and N-acetyl-alanine were increased. Glutamate dysregulation has previously been associated with heat stroke (Chauhan et al., 2017; He et al., 2023; Niu et al., 2003; Yang et al., 1998). In males, two metabolites, N-acetyl-phenylalanine and ornithine, had MDA scores > 0.01 (**Figure 7C**). Like ornithine, the levels of N-acetyl-phenylalanine decreased after heat exposure compared to control male mice. The identification of ornithine in both the heatmap and random forest analysis suggests it is a robust marker of heat-induced metabolic changes. A decrease in N-acetyl-phenylalanine and ornithine may indicate a disruption in amino acid metabolism, impaired protein synthesis, and dysfunction in the urea cycle affecting nitrogen metabolism and ammonia detoxification (Cui et al., 2022; Fox et al., 2019; Zhao et al., 2021). Heat exposure has been linked to kidney disease (Chapman et al., 2021), which in turn can affect the urea cycle (Ippolito et al., 2014). In contrast to our findings, the earlier work reported that phenylalanine and ornithine levels increased after 24 and 48 hours of heat exposure (Ippolito et al., 2014). These differences may be due to the different heat exposure paradigms and the potential progression to renal dysfunction in the previous study.

**Figure 6.**
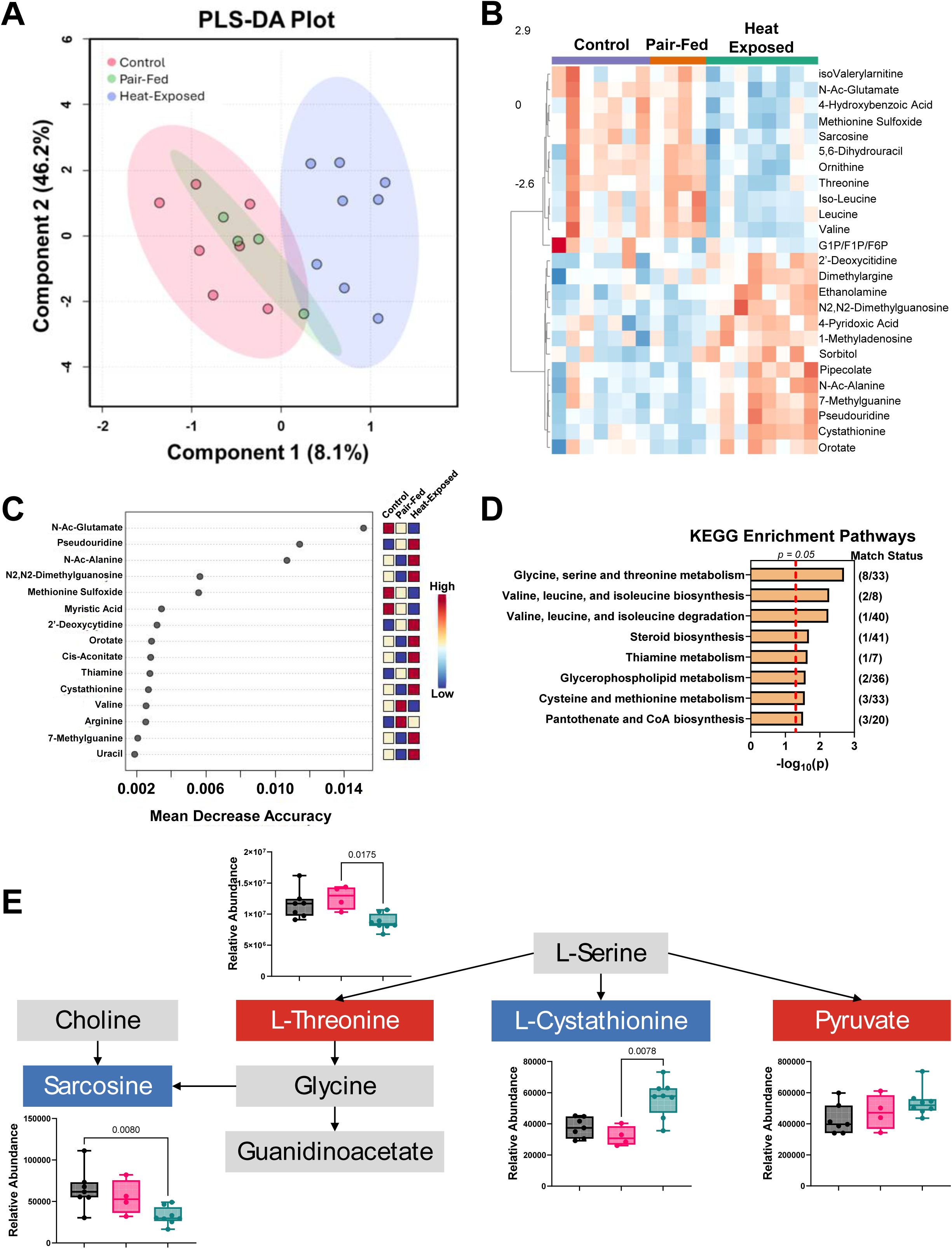

#### 3.5.2. Analysis of Metabolic KEGG pathways

KEGG enrichment analysis was performed using the altered serum metabolites to assess biological pathway-level changes. Heat exposure significantly impacted eight KEGG pathways in female mice (**Figure 6D**) and 20 pathways in male mice (**Figure 7D**), indicating more pronounced metabolic effects in males. Among the enriched pathways, “Glycine, Serine, and Threonine Metabolism” was significantly affected by heat exposure in both sexes (**Figures 6E** and **7E**). This pathway is involved in carbohydrate and energy utilization efficiency (Wu et al., 2021). Within this pathway, pyruvate levels increased while sarcosine levels decreased in both sexes. However, L-threonine and L-cystathionine were altered in opposite directions between females and males. Choline, glycine, and guanidinoacetate were elevated in males after heat exposure but remained unchanged in females. Interestingly, a previous study has shown that a 3-day heat exposure decreased choline levels and associated these levels with increased anxiety (Fang et al., 2023). Moreover, the elevated guanidinoacetate levels may reflect disruptions in creatine synthesis or guanidinoacetate methyltransferase (GAMT), which can lead to neurocognitive impairments, as demonstrated in rodent models (Hanna-El-Daher et al., 2015). These metabolic shifts align with reports that heat-related illnesses can cause neurological complications, including cognitive impairment (Yoneda et al., 2024).

**Figure 7.**
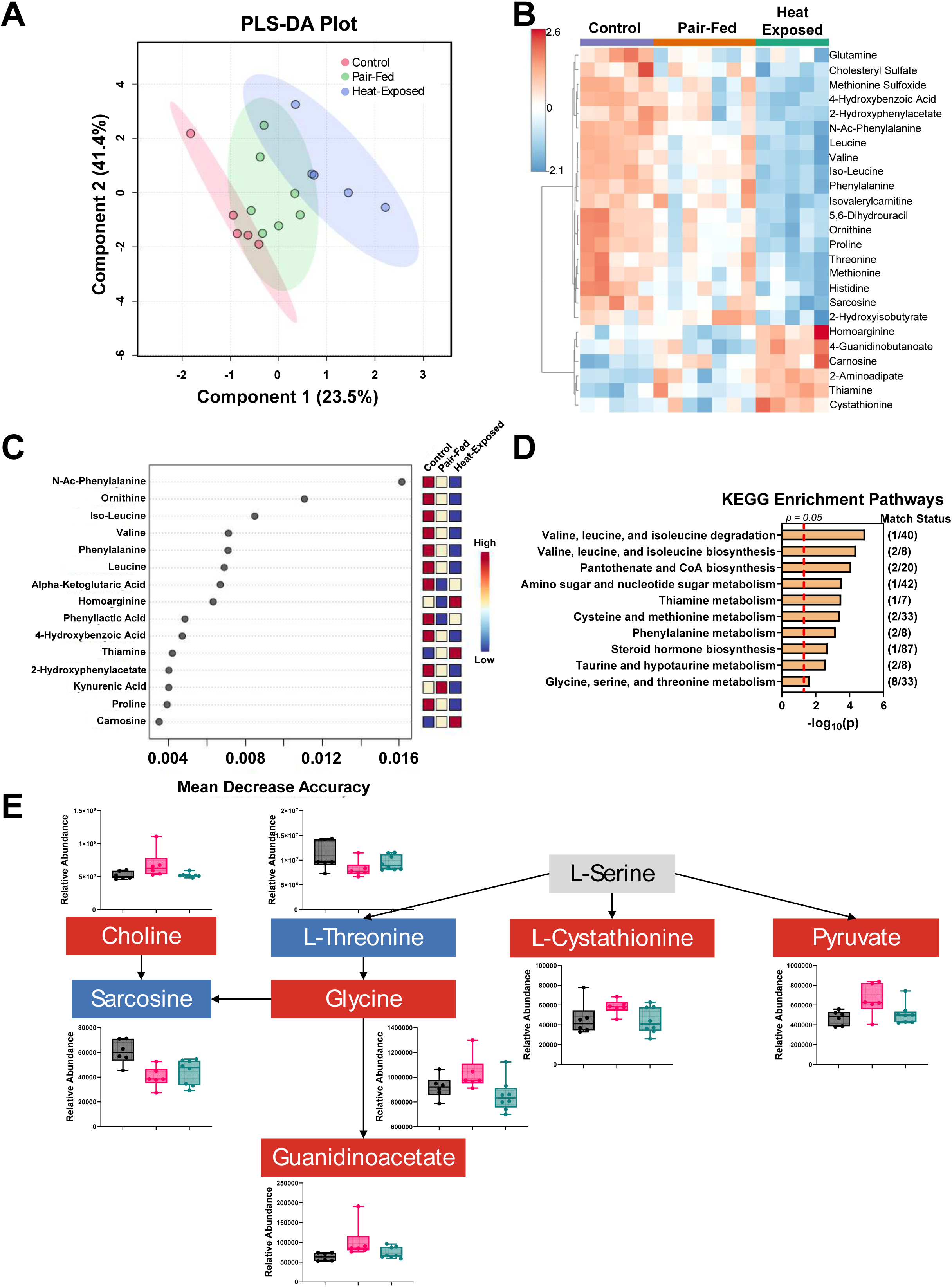

### 3.6. Correlation analysis of cytokines, gene expression and metabolomics data

Correlation-based network analyses were performed by integrating lung transcriptomic profiles with serum metabolites, serum cytokines, and lung cytokines in both male and female mice to characterize the systems-level impact of heat exposure further. These analyses focused on strong correlations (R > 0.9, p < 0.05) to highlight key immunometabolic interactions (**Figure 8**). These networks were constructed for three group comparisons: (1) heat-exposed vs. control, (2) heat-exposed vs. pair-fed, and (3) pair-fed vs. control. This approach enabled us to differentiate changes resulting from heat exposure from those associated with reduced feed intake and characterize sex-dependent systemic and pulmonary immunometabolic signatures altered in response to heat exposure.

**Figure 8.**
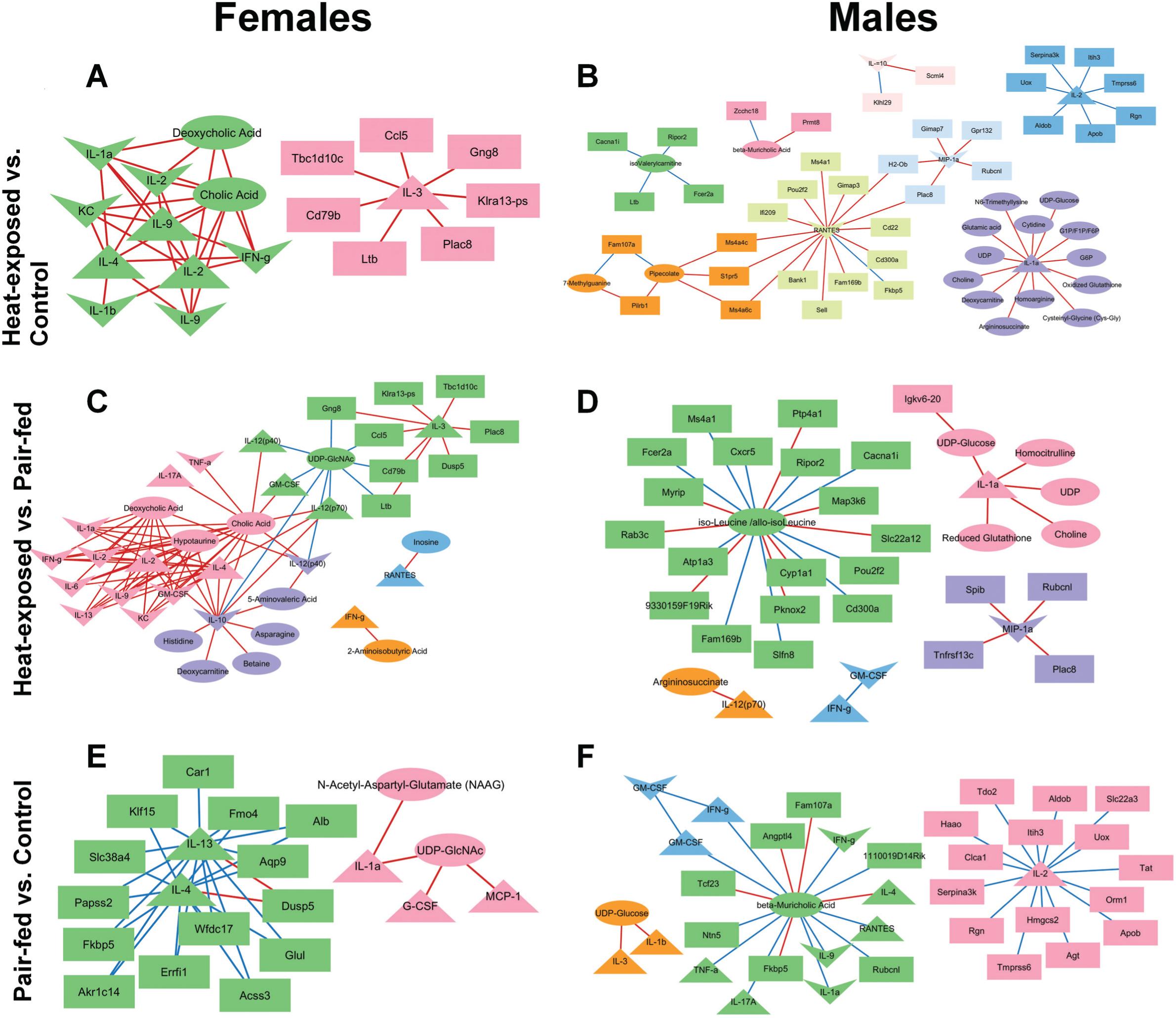

#### 3.6.1. Heat-exposed and control correlation network analysis

In female mice, the correlation network consisted of two distinct clusters, comprising seven DEGs, two endogenous metabolites, four serum cytokines, and six lung cytokines (**Figure 8A**). All 36 observed correlations were positive. Lung IL-1α was positively correlated with serum IL-4, IL-9, and IL-2 as well as with the bile acids cholic acid and deoxycholic acid. These findings suggest a link between inflammatory signaling and altered bile acid homeostasis in heat-exposed female mice (Fleishman & Kumar, 2024; Oleszycka et al., 2023). The networks in male mice were more complex, comprising eight clusters that contained 35 DEGs, 17 metabolites, two serum cytokines, and three lung cytokines, with 39 positive and 16 negative correlations (**Figure 8B**). Three cytokines elevated after heat exposure, including lung MIP-1α, serum RANTES, and serum IL-1α (**Figure 2**), were central hubs within the network. Lung MIP-1α was positively correlated with five DEGs, lung RANTES was positively associated with 14 DEGs, and serum IL-1α was positively correlated with 13 endogenous metabolites. *Fcer2a*, which was upregulated after heat exposure (**Figure S3A**), showed a positive correlation with isovalerylcarnitine. These associations suggest that specific genes play a central role in orchestrating immunometabolic responses in heat exposed male mice (Huang et al., 1992; Lan et al., 2008; Menten et al., 2002).

Two DEGs, *Plac8* (Placenta Associated 8) and *Ltb* (Lymphotoxin beta), were identified in both female and male correlation networks. Both genes were significantly upregulated in response to heat exposure. In females, *Plac8* and *Ltb* were positively correlated with serum IL-3. In males, *Plac8* was positively correlated with lung MIP-1α and RANTES, while *Ltb* was negatively correlated with isovalerylcarnitine. Increased *Plac8* is linked with Th1/Th2/Th17/Treg balance disruption in mice (Yang et al., 2021) and monocyte function modulation in humans (Zhang et al., 2024). Elevated *Ltb* is linked to pulmonary inflammation in mice (Miki et al., 2023) and regulates the function and movement of T lymphocytes in mice (Alfituri et al., 2020; Kuprash et al., 1996; Zhao et al., 2014). The consistent correlations between *Plac8* and *Ltb* with both serum and lung cytokines support our findings that acute heat exposure provokes a coordinated inflammatory response in both female and male mice.

#### 3.6.2. Heat-exposed and pair-fed correlation network analysis

In female mice, the heat-exposed vs. pair-fed correlation network comprised five clusters containing eight DEGs, 11 serum metabolites, nine serum cytokines, and 11 lung cytokines, with a total of 72 positive and nine negative correlations (**Figure 8C**). Among the central hubs were IL-12(p40) and RANTES, both of which were elevated in serum following heat exposure compared to pair-fed animals (**Figure 2**). Lung and serum IL-12(p40) were positively correlated with two bile acids, cholic acid and deoxycholic acid, and 5-aminovaleric acid, and negatively correlated with uridine diphosphate N-acetylglucosamine (UDP-GlcNAc). In male mice, the network consisted of five clusters, comprising 23 DEGs, seven metabolites, three serum cytokines, and two lung cytokines, with 19 positive and 11 negative correlations (**Figure 8D**). Two cytokines elevated in heat-exposed males, lung MIP-1α and serum IL-1α (**Figure 2**), were key network hubs. Lung MIP-1α was positively correlated with four DEGs, while serum IL-1α was positively associated with five metabolites. These results are consistent with immunometabolic responses following heat exposure in both sexes (Eom et al., 2022; Wang & Medzhitov, 2019).

When comparing networks between sexes, *Plac8* and lung GM-CSF appeared in both female and male heat-exposed vs. pair-fed networks (**Figures 8C** and **8D**) *Plac8* was a significantly upregulated hub gene in both sexes. In females, *Plac8* was positively correlated with serum IL-3, consistent with findings in the heat-exposed vs. control comparison (**Figure 8A**). In males, *Plac8* was positively correlated with lung MIP-1α, again mirroring the prior comparison (**Figure 8B**). Lung GM-CSF, while not significantly altered at the group level, was positively correlated with cholic acid and negatively correlated with UDP-GlcNAc in females. In contrast, GM-CSF levels in males were negatively correlated with interferon-γ (IFN-γ). These divergent associations suggest sex-specific immunometabolic regulatory roles for GM-CSF, a pleiotropic cytokine that modulates myeloid cell function and T-cell activation (Bhattacharya et al., 2015). Together with the networks of heat-exposed vs. control mice (**Figures 8A** and **8B**), the positive correlations between *Plac8* and cytokines in lung and serum support an inflammatory response to heat exposure in male and female mice (Yang et al., 2021; Zhang et al., 2024).

#### 3.6.3. Pair-fed and control correlation network analysis

In female mice, the pair-fed vs. control network included 14 DEGs, two metabolites, and five serum cytokines, forming two clusters with six positive and 25 negative correlations (**Figure 8E**). Serum IL-1α, reduced in pair-fed mice (**Figure 2D**), was positively correlated with N-acetyl-aspartyl-glutamate and UDP-GlcNAc, suggesting that even modest reductions in feed intake can influence systemic inflammation (Kvidera et al., 2017; Stumpf et al., 2023). In males, the network analysis identified 22 DEGs, two metabolites, nine serum cytokines, and four lung cytokines across four clusters, with seven positive and 28 negative correlations (**Figure 8F**). Serum IL-17α, lung IL-1α, and serum RANTES were negatively correlated with β-muricholic acid, a bile acid that regulates metabolic and immune pathways via FXR signaling (Chávez-Talavera et al., 2017; Sayin et al., 2013).

IL-4 appeared in both sexes. In females, it was negatively correlated with several metabolism- and inflammation-related DEGs, while in males, it was positively correlated with β-muricholic acid. Not surprisingly, many of the genes identified in the network analysis were related to metabolism (Abdollahi et al., 2023; Bayram et al., 2022; Brylski et al., 2022; Phillips & Shephard, 2019; Rižner & Penning, 2014; Sasse et al., 2013; Wang et al., 2024a) and inflammation (Kasmani et al., 2023; Russo et al., 2019; Seo et al., 2017; Tesse et al., 2024; Zou & Zeng, 2023). These patterns suggest that nutrient restriction alone can modestly alter immunometabolic networks through IL-4 and bile acid signaling.

## 4. Conclusions

Eight hours of gradual extreme heat exposure in mice led to coordinated, sex-specific physiological, immunological, metabolic, and transcriptomic changes. Although body weight and feed consumption were not significantly affected, water intake increased, reflecting a thermoregulatory response. Unlike prior studies using acute stress models, our gradual heat paradigm revealed a trend toward increased feeding, suggesting different behavioral adaptations depending on the type of heat exposure (e.g., sudden vs. gradual). Cytokine profiling revealed a complex immune response: the reduced levels of IL-1α and IL-17α suggest suppressed acute inflammation and mucosal defense, while elevated levels of IL-12(p40), RANTES, MIP-1α, and MIP-1β indicated enhanced immune cell recruitment that could lead to chronic inflammation if repeated exposure to heat occurs, or inflammation does not resolve. Serum metabolomics showed consistent reductions in several metabolites across sexes, including amino acids and bile acid-related compounds, with enrichment of pathways such as “Glycine, Serine, and Threonine Metabolism,” particularly in males. These findings reflect stress-induced metabolic reprogramming and potential impairments in nitrogen balance and energy utilization. Although the alpha and beta diversity of the gut microbiome remained unchanged, random forest analysis identified taxa, including reduced Lactobacillus in males, that are potentially relevant to immune modulation and mucosal health. Network analyses integrating transcriptomics, metabolites, and cytokines revealed *Plac8* and *Ltb* as conserved immune-regulatory hubs correlated with cytokines and bile acid metabolites. These results underscore the bidirectional relationship between immune signaling and metabolism during heat stress. Sex-specific differences in molecular network structure further suggest that male and female mice utilize distinct strategies to cope with thermal stress. Collectively, this study advances our systems-level understanding of how gradual extreme heat stress reshapes immune and metabolic function, offering insight into mechanisms that may underlie chronic inflammation and metabolic dysregulation in response to extreme heat events.

## Acknowledgments

This work was supported by grants NIH P30 ES005605 and T32HL166134 from the National Institutes of Environmental Health Science and National Institutes of Health (National Heart, Lung, and Blood Institute (NHLBI)). The authors thank Darby Forsyth from the University of Iowa for her assistance with animal exposures and tissue and sample collection. We also thank Dr. Hui Wang, Dr. Amanda J. Bullert, and Nicole Breese from the University of Iowa for their contributions to tissue and sample collection. We acknowledge the support provided by the FOEDRC Metabolic Phenotyping Core Facility at the University of Iowa for animal exposures, the University of Iowa Microbiome Core for providing 16S rRNA sequencing services, and the Northwest Metabolomics Research Center (NM-MRC) for providing metabolomics services.

## Declaration of generative AI and AI-assisted technologies in the writing process

During the preparation of this work, the author(s) used ChatGPT (chat.openai.com), Grammarly (grammarly.com), and Copilot (copilot.microsoft.com) to edit sections of the paper. After using these tools/services, the authors reviewed and edited the content as needed and take full responsibility for the content of the published article.

## CRediT Authorship Statement

**Laura E. Dean**: Conceptualization, Validation, Formal analysis, Data curation, Writing – original draft, Visualization; **Andrea Adamcakova-Dodd**: Conceptualization, Investigation, Resources, Writing – review & editing; **Hans-Joachim Lehmler**: Conceptualization, Methodology, Validation, Writing – review & editing, Supervision, Project administration, Funding acquisition on of Competing Interests

## Research Data

All data reported in this manuscript are publicly available on Iowa Research Online and can be accessed at DOI: 10.25820/data.007638

