## Supplementary Material for "Ramping up the Heat: Induction of Systemic and Pulmonary Immune Responses and Metabolic Adaptations in Mice"

**Corresponding Author:**	Dr. Hans-Joachim Lehmler

Iowa City, IA 52242-5000

**E-Mail Addresses:** (L. E. Dean), (A. Adamcakova-Dodd), and (H. Lehmler)

**Table of Contents**

| **Table S1.** Concentrations of cytokines and chemokines (pg/mL); mean ± standard deviation | S3 |
| --- | --- |
| **Table S2.** Body (g), feed (g), water (g), and relative organ weights (organ/body weights %); mean ± standard deviation | S5 |
| **Figure S1.** Alpha Diversity was not altered in (A) female or (B) male mouse microbiomes after heat exposure and pair-feeding. Beta Diversity was not changed in (C) female or (D) male mouse microbiomes after heat exposure and pair-feeding. Alpha diversity was measured by Shannon Diversity at the feature level. PCoA and Jensen-Shannon Divergence at the feature level were used to measure beta diversity. | S6 |
| **Figure S2.** PCA plots showed no separation in transcriptomic profiles in the hippocampus (A) of female or (B) male mice, or in the lung (C) of female or (D) male mice, after heat exposure or pair-feeding. | S7 |
| **Figure S3.** Expression patterns of differentially expressed genes (DEGs) shared across altered biological processes between female and male heat-exposed mice and their control counterparts are shown in panels (A-D). DEG normalized counts are presented as mean ± standard deviation. | S8 |

**Table S1.** Concentrations of cytokines and chemokines (pg/mL); mean ± standard deviation

| **Cytokine**  **or**  **Chemokine** | **Sex** | **Lung Homogenate** | | | **Serum** | | |
| --- | --- | --- | --- | --- | --- | --- | --- |
|  |  | **Control**  **(n=9M, 8F)** | **Exposed**  **(n=9M, 8F)** | **Pair-Fed**  **(n=6M, 6F)** | **Control**  **(n=8M, 7F)** | **Exposed**  **(n=8M, 6F)** | **Pair-Fed**  **(n=8M, 8F)** |
| **Eotaxin** | M | 1591.9±550.9 | 1764.9±245.9 | 1538.7±144.9 | 1509.7±656.5 | 1676.1±644.2 | 1141.6±316.2 |
|  | F | 1474.6±250.1 | 1391.2±480.1 | 1269.6±242.5 | 2058.7±625.7 | 2189.9±1124.0 | 2409.6±784.0 |
| **G-CSF** | M | 1.1±0.2 | 1.1±0.4 | 1.6±0.2 | 58.7±38.2 | 67.9±41.1 | 38.4±7.2 |
|  | F | 1.3±0.5 | 1.3±0.6 | 1.0±0.3 | 32.8±9.8 | 31.2±10.6 | 34.5±15.6 |
| **GM-CSF** | M | 11.5±10.1 | 7.4±9.2 | 16.7±10.4 | 3.2±1.3 | 11.2±8.9 | 6.5±4.4 |
|  | F | 16.6±11.7 | 17.8±20.2 | 9.8±11.0 | 15.1±10.6 | 9.8±8.0 | 6.4±6.7 |
| **IFN-γ** | M | 34.9±13.2 | 39.1±8.0 | 30.7±7.9 | 2.4±1.2 | 3.7±3.7 | 2.3±2.3 |
|  | F | 34.6±8.8 | 51.5±34.8 | 32.8±2.7 | 5.1±3.5 | 6.0±3.0 | 3.1±3.0 |
| **IL-1α** | M | 21.7±7.5 | 24.4±4.6 | 20.2±6.2 | 76.3±74.1 | 22.0±15.4 | 16.7±13.4 |
|  | F | 23.6±5.9 | 35.9±31.0 | 20.6±3.4 | 15.8±6.1 | 32.4±36.6 | 21.2±12.3 |
| **IL-1β** | M | 4.5±0.9 | 4.9±1.0 | 4.7±1.4 | 3.1±6.2 | 1.2±0.6 | 1.3±0.4 |
|  | F | 5.1±0.4 | 5.8±2.0 | 4.9±1.4 | 1.2±0.4 | 0.8±0.5 | 0.9±1.2 |
| **IL-2** | M | 45.8±14.5 | 50.8±10.5 | 45.4±16.4 | 1.0±0.0 | 1.1±0.1 | 0.9±0.4 |
|  | F | 49.6±9.6 | 54.5±6.7 | 45.9±4.9 | 1.0±0.2 | 1.0±0.0 | 1.0±0.2 |
| **IL-3** | M | 2.6±1.7 | 2.3±1.9 | 4.3±2.0 | 0.4±0.2 | 3.4±7.7 | 0.4±0.1 |
|  | F | 3.3±1.7 | 2.4±2.4 | 1.9±1.6 | 0.5±0.3 | 0.8±0.5 | 0.9±1.0 |
| **IL-4** | M | 1.4±1.1 | 1.7±0.9 | 1.6±0.7 | 0.4±0.2 | 0.8±0.8 | 0.5±0.0 |
|  | F | 1.4±0.7 | 2.2±1.5 | 1.2±0.8 | 0.2±0.3 | 0.5±0.6 | 0.8±0.5 |
| **IL-5** | M | 10.4±4.5 | 9.4±3.2 | 8.9±4.0 | 1.5±1.0 | 3.0±2.4 | 2.9±1.5 |
|  | F | 8.6±3.4 | 9.8±2.2 | 10.1±2.7 | 2.6±1.9 | 2.7±1.3 | 2.4±2.0 |
| **IL-6** | M | 2.6±2.0 | 3.0±1.1 | 2.9±1.1 | 2.5±0.0 | 2.5±0.0 | 2.5±0.0 |
|  | F | 2.3±2.4 | 5.4±4.9 | 1.8±1.2 | 2.5±0.0 | 3.8±3.0 | 2.5±0.0 |
| **IL-9** | M | 44.2±21.1 | 53.6±14.9 | 34.3±16.3 | 1.7±0.6 | 2.7±2.0 | 3.8±4.5 |
|  | F | 45.5±18.6 | 75.5±61.5 | 41.1±7.7 | 3.2±2.6 | 5.3±7.9 | 3.3±2.6 |
| **IL-10** | M | 1.3±2.8 | 0.3±0.9 | 1.4±1.7 | 2.2±1.0 | 3.0±2.1 | 2.4±1.1 |
|  | F | 0.9±2.5 | 4.9±10.1 | 0.7±1.7 | 3.6±2.5 | 3.3±2.5 | 3.4±1.8 |
| **IL-12(p40)** | M | <13.7 | <13.7 | <13.7 | 261.8±116.6 | 405.4±154.6 | 232.8±73.1 |
|  | F | <13.7 | <13.7 | <13.7 | 262.6±54.1 | 374.1±75.3 | 374.1±161.3 |
| **IL-12(p70)** | M | <4.7 | <4.7 | <4.7 | 5.2±4.9 | 3.3±0.9 | 3.3±0.0 |
|  | F | <4.7 | <4.7 | <4.7 | 5.3±5.2 | 5.8±5.9 | 5.2±5.2 |
| **IL-13** | M | 16.9±16.3 | 11.9±11.7 | 7.7±15.5 | 8.4±0.5 | 7.6±1.8 | 8.5±0.0 |
|  | F | 22.8±21.8 | 39.3±42.1 | 23.0±10.4 | 7.5±2.8 | 8.1±2.5 | 7.9±1.9 |
| **IL-17α** | M | 2.1±1.2 | 1.5±0.5 | 3.2±1.7 | 17.1±11.8 | 24.4±20.2 | 16.7±9.6 |
|  | F | 2.4±1.1 | 1.9±0.7 | 1.8±1.2 | 25.9±20.9 | 20.7±11.8 | 11.0±18.0 |
| **KC** | M | 37.7±9.6 | 38.2±4.2 | 39.1±7.3 | 31.8±15.5 | 33.5±16.5 | 27.0±7.7 |
|  | F | 39.7±5.5 | 43.4±20.7 | 35.7±5.4 | 35.4±8.9 | 42.2±21.2 | 41.7±7.5 |
| **MCP-1** | M | 205.8±50.2 | 215.6±43.8 | 232.5±44.2 | 187.8±384.6 | 68.5±39.3 | 47.2±23.9 |
|  | F | 236.7±40.7 | 267.0±65.8 | 205.1±40.6 | 54.2±18.0 | 62.0±26.2 | 48.0±24.1 |
| **MIP-1α** | M | 23.9±4.1 | 33.1±8.1 | 23.3±4.5 | 0.4±0.0 | 0.5±0.2 | 0.4±0.0 |
|  | F | 30.2±9.7 | 43.9±18.2 | 27.7±10.1 | 0.4±0.1 | 0.6±0.4 | 0.4±0.2 |
| **MIP-1β** | M | 118.2±27.3 | 142.7±36.5 | 129.3±32.5 | 7.4±0.0 | 21.0±20.8 | 15.6±20.2 |
|  | F | 134.4±45.5 | 167.9±43.8 | 114.0±32.9 | 16.9±18.7 | 25.3±26.9 | 15.7±17.6 |
| **RANTES** | M | 614.1±129.8 | 833.7±204.0 | 597.3±151.5 | 21.9±14.7 | 23.2±13.7 | 7.3±5.7 |
|  | F | 922.7±684.7 | 1017.8±278.4 | 743.4±314.3 | 25.7±10.1 | 44.6±14.3 | 24.6±14.6 |
| **TNF-α** | M | 119.5±43.5 | 122.5±37.2 | 123.0±31.9 | 5.8±3.4 | 11.9±14.1 | 4.9±6.5 |
|  | F | 134.3±31.3 | 157.0±54.1 | 120.4±22.2 | 7.5±4.9 | 6.2±4.4 | 5.0±5.2 |

**Table S2.** Body (g), feed (g), water (g), and relative organ weights (organ/body weights %); mean ± standard deviation

|  | **Sex** | **Control**  **(n=9M, 8F)** | **Exposed**  **(n=9M, 8F)** | **Pair-Fed**  **(n=6M, 6F)** |
| --- | --- | --- | --- | --- |
| **Starting Weight** | M | 25.0±0.7 | 25.2±1.6 | 26.5±0.8 |
|  | F | 20.3±1.3 | 20.7±1.1 | 20.2±1.8 |
| **Ending Weight** | M | 24.8±0.6 | 25.3±1.8 | 26.1±0.9 |
|  | F | 20.1±1.5 | 20.4±1.2 | 19.8±1.6 |
| **Feed Consumption** | M | 0.7±0.3 | 0.8±0.3 | 0.6±0.3 |
|  | F | 0.8±0.3 | 0.8±0.3 | 0.5±0.2 |
| **Water Consumption** | M | 0.8±0.3 | 4.8±2.4 | 1.9±1.9 |
|  | F | 0.9±0.4 | 3.7±0.5 | 2.0±1.2 |
| **Brain** | M | 1.8±0.1 | 1.7±0.1 | 1.7±0.1 |
|  | F | 2.2±0.1 | 2.2±0.1 | 2.2±0.1 |
| **Lung** | M | 0.7±0.1 | 0.7±0.1 | 0.7±0.1 |
|  | F | 0.8±0.2 | 0.8±0.3 | 0.8±0.2 |
| **Liver** | M | 4.3±0.2 | 3.9±0.1 | 3.9±0.3 |
|  | F | 4.8±0.6 | 4.5±0.7 | 4.2±0.5 |
| **Combined Kidneys** | M | 1.3±0.1 | 1.3±0.1 | 1.3±0.1 |
|  | F | 1.4±0.3 | 1.4±0.2 | 1.4±0.2 |
| **Thymus** | M | 0.2±0.0 | 0.1±0.0 | 0.2±0.0 |
|  | F | 0.2±0.1 | 0.2±0.1 | 0.2±0.0 |
| **Spleen** | M | 0.3±0.0 | 0.3±0.0 | 0.3±0.0 |
|  | F | 0.4±0.1 | 0.5±0.1 | 0.4±0.1 |
| **Heart** | M | 0.5±0.1 | 0.6±0.1 | 0.6±0.1 |
|  | F | 0.6±0.1 | 0.6±0.1 | 0.6±0.1 |
| **Combined Testes/Ovary** | M | 0.8±0.0 | 0.9±0.3 | 0.8±0.0 |
|  | F | 0.4±0.4 | 0.4±0.4 | 0.4±0.4 |

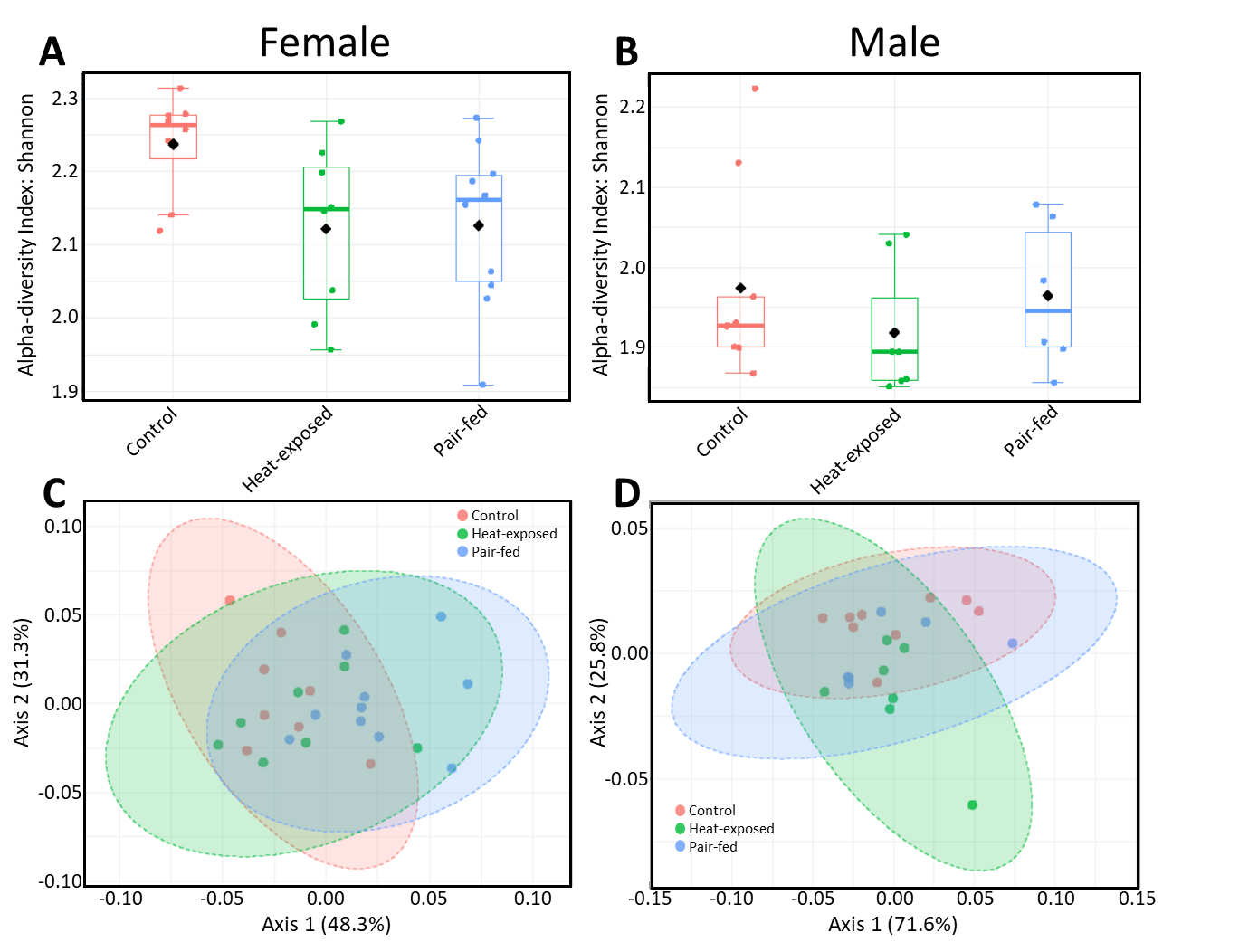

**Figure S1.** Alpha Diversity was not altered in (A) female or (B) male mouse microbiomes after heat exposure and pair-feeding. Beta Diversity was not changed in (C) female or (D) male mouse microbiomes after heat exposure and pair-feeding. Alpha diversity was measured by Shannon Diversity at the feature level. PCoA and Jensen-Shannon Divergence at the feature level were used to measure beta diversity.

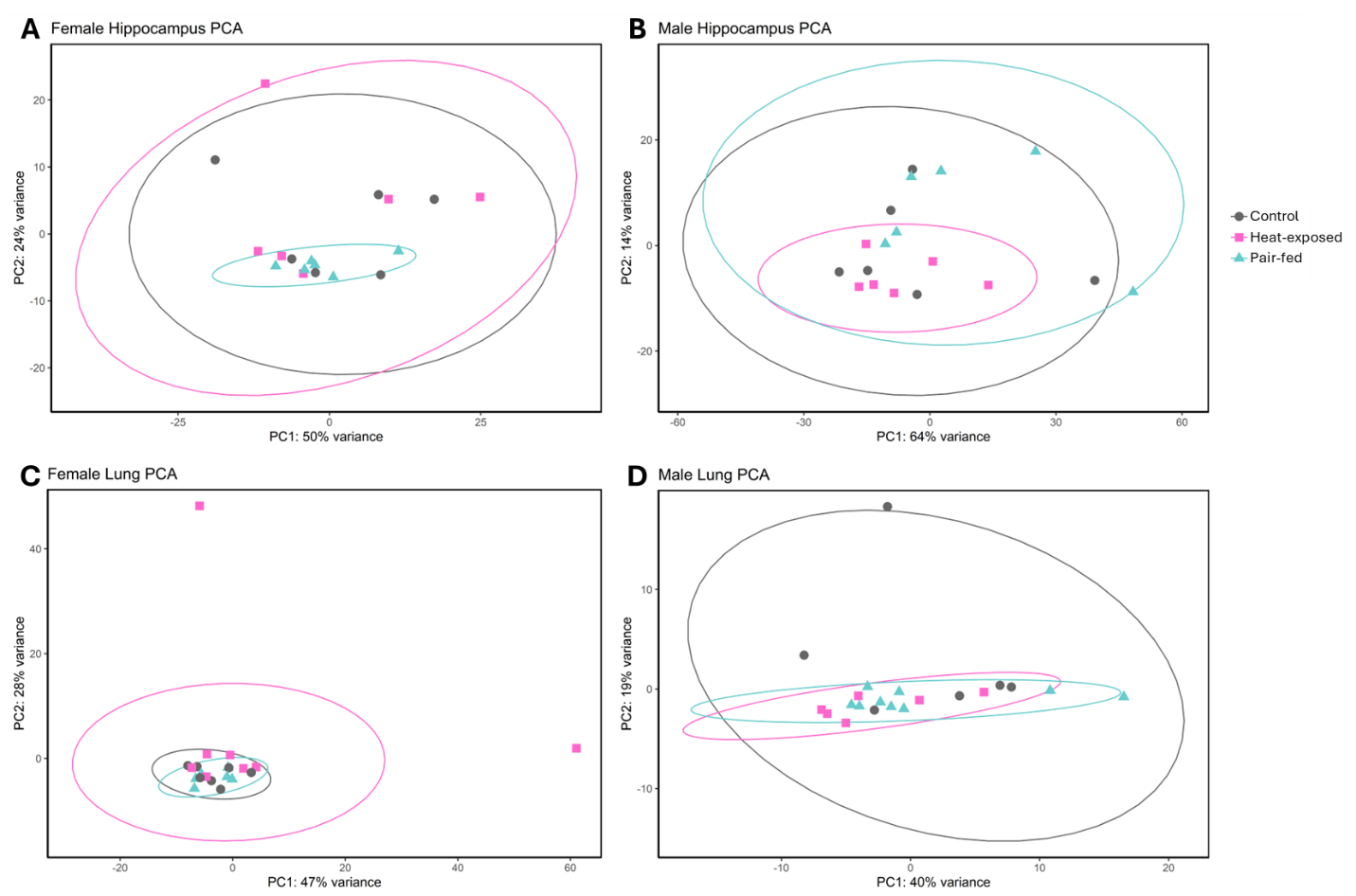

**Figure S2.** PCA plots showed no separation in transcriptomic profiles in the hippocampus (A) of female or (B) male mice, or in the lung (C) of female or (D) male mice, after heat exposure or pair-feeding.

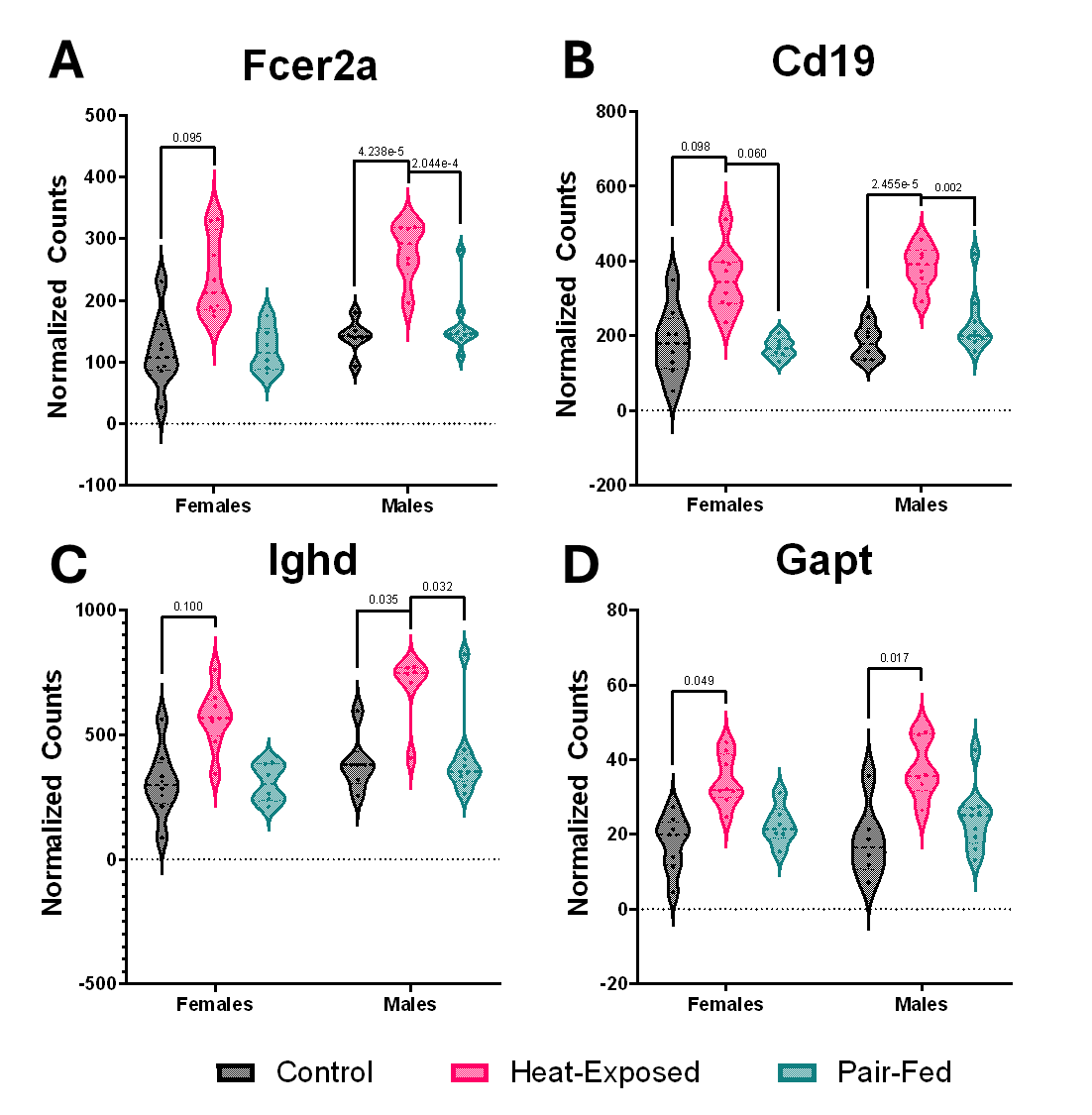

**Figure S3.** Expression patterns of differentially expressed genes (DEGs) shared across altered biological processes between female and male heat-exposed mice and their control counterparts are shown in panels (A-D). DEG normalized counts are presented as mean ± standard deviation.
